# Interference between flexible and adaptive reaching control

**DOI:** 10.1101/2025.06.23.660708

**Authors:** Astrid Doyen, Philippe Lefèvre, Frédéric Crevecoeur

## Abstract

Humans rapidly update the control of an ongoing movement following changes in contextual parameters. This involves adjusting the controller to exploit redundancy in the movement goal, such as when reaching for a narrow or wide target, and adapting to dynamic changes such as velocity-dependent force fields (FFs). Although flexible control and motor adaptation are computationally distinct, the fact that both unfold within the same movement suggests they may share common neural resources for task-specific adjustments. To test this hypothesis, we conducted a series of experiments combining changes in the target structure and a force field presented separately or in combination. Seventy-six human participants (both sexes) took part in this study, with each experiment involving different participants. They were asked to reach for a target that could change from a narrow square to a wide rectangle between or during trials. Step loads were used to assess whether participants exploited target redundancy. In a separate experiment, we added a force field in addition to target changes and step loads. Our results revealed a reduced ability to exploit target redundancy when sudden target changes occurred concurrently with FF adaptation. Furthermore, the magnitude of adaptation was reduced when step loads were added to the FF. Crucially, this interference emerged specifically when all perturbations impacted motor execution simultaneously. These results indicate that flexible control and motor adaptation interact in a non-trivial manner, suggesting possible overlap between their underlying neural mechanisms, and a clear identification of the timescale at which they are engaged – namely, during movement.

**Significant statement:** Humans rapidly adapt to changes in task demands, such as target structure changes or exposure to force fields (FFs). These two types of adjustments occur within a single movement, suggesting potential interactions between them. Our experiments revealed that the combination of FF exposure with online target shape changes selectively reduced participants’ ability to exploit target redundancy, while the combination of FF and step loads led to a reduced extent of motor adaptation. These findings confirm that motor adaptation occurs not only between trials but also during movement. The selective nature of the observed interference highlights an interplay between flexible control and motor adaptation, underscoring the importance of understanding the timing of these processes to better characterise their underlying neural circuits.

## Introduction

Reaching movements are fundamental daily actions that sometimes require flexibility to cope with external perturbations like forces applied to the hand or changes in target shape or position. These factors impact planning and execution strategies, as evidenced by the effects of visual and mechanical perturbations on movement kinematics and feedback control strategies (Franklin and Wolpert, 2008; Izawa and Shadmehr, 2008; Kurtzer et al., 2008; Crevecoeur et al., 2013).

Several studies have explored humans’ ability to select control policies based on task demands by designing experiments where participants have flexibility associated with a state variable (position or velocity). For instance, humans select different control policies for shooting or stopping at a target (Liu & Todorov, 2007; Česonis & Franklin, 2022). Similarly, reaching behaviour differs depending on target shape, with greater endpoint dispersion for wide bars than dots (Nashed et al., 2012). This supports the minimal intervention principle (Todorov and Jordan, 2002a), which states that deviations from trajectory are corrected only if they interfere with the task goals. De Comite et al. (2021) further showed that such flexibility in corrections occurred even when the target changed during movement, highlighting control policy updates during movement.

In parallel, humans also adapt to new dynamic environments to recover the intended performance when exposed to a predictable perturbation. For instance, under a force field (FF) that deviates the hand from the straight path, participants gradually reduce the deviation, restoring straighter movements (Shadmehr and Mussa-Ivaldi, 1994; Dizio and Lackner, 1995; Singh and Scott, 2003; Lackner and DiZio, 2005). Feedback mechanisms play a central role in motor adaptation, with the brain adjusting a task-specific feedback controller to the requirements of novel motor skills (Ahmadi-Pajouh et al., 2012; Cluff and Scott, 2013; Joiner et al., 2017; Maeda et al., 2020). In addition to controller adjustments between movements, a fast timescale for motor adaptation is also present, influencing the execution of a single movement when different FFs are presented randomly to the participants (Crevecoeur et al., 2020b).

Much of the experimental work about reaching movements has been guided by optimal feedback control theory, which posits that task goals and environmental dynamics shape efficient movement strategies. In this framework, an optimal goal-dependent control strategy determines feedback gains during movement based on task constraints and limb dynamics (Todorov and Jordan, 2002b; Scott, 2004; Liu and Todorov, 2007). We focus on two types of control changes: those related to task demands and those related to changes in dynamics. Mathematically, task demands are expressed through the cost function, representing the goals and constraints of the task. For example, when the goal target switches during movement, penalties on behavioural performance are affected. In contrast, changes in dynamics refer to adjustments in internal models representing limb or environment dynamics. Although these operations are theoretically distinct, both changes in movement due to task updates and adaptation to changes in dynamics occur within ∼250ms following a contextual perturbation (Kalidindi and Crevecoeur, 2023). Considering a typical movement time of ∼500ms, this similarity in timing suggests that both controller updates and adaptation may occur within a reaching movement.

To explore their potential overlap, we examined behavioural interference during a motor task with changes in goal target structure and step loads (‘flexible control’) with or without FF (‘adaptation’). Two hypotheses were formulated: if participants performed similarly in separated and combined subtasks, it would suggest independent neural circuits; conversely, behavioural differences would indicate interference. Consistent with the latter hypothesis, we found differences in participants’ ability to exploit target redundancy when exposed to the FF, but only when the target switch occurred during movement. Moreover, motor adaptation was reduced when step loads were applied during movement. Importantly, these interferences were not uniform across all conditions but emerged selectively when both adaptive and flexible control mechanisms were recruited simultaneously. These results indicate the interdependence of neural circuits involved in the online modification of the control policy and motor adaptation.

## Materials and methods

### Participants

This study involved three groups of participants. The first group of 20 participants (18 right-handed and 2 left-handed, 11 females), aged from 20 to 31 was assigned to main Experiment 1. The second group, composed of 20 right-handed participants (11 females) with ages ranging from 21 to 26, performed main Experiment 2. Note that these two experiments involved different participants, hence we chose to refer to these datasets as two different experiments. For the control experiments, 36 participants (28 females) were involved, with ages ranging from 18 to 32, and divided into 3 groups of 12 participants, one for each control experiment. All participants were unaware of the purpose of the study, had normal or corrected to normal vision, and had no known neurological or motor disorder. The procedures were approved by the ethics committee at the host institution (Comité d’Éthique Hospitalo-Facultaire, UCLouvain) and participants received financial compensation for their time.

### Setup

All experiments were conducted using the KINARM endpoint robotic device (KINARM, Kingston, ON, Canada). Participants were seated on an adjustable chair, facing the robotic device. They were instructed to grasp the handle of the right robotic arm using their right arm. This robotic arm allowed movements in the horizontal plane. They were seated at the beginning of each trial in such a way that their elbow and shoulder lie in a plane orthogonal to the movement plane as can be seen in Figure 1A. The direct view of both the participant’s hand and the robotic arm was blocked. An augmented-reality display was used to project the hand-aligned cursor (radius 0.5cm), the virtual targets (Figure 1A) and a score corresponding to the sum of all successful trials to keep participants engaged in the task. Muscular activity of two muscles of interest (Pectoralis Major and Posterior Deltoid) was measured using surface electrodes (Bagnoli surface EMG Sensor, Delsys Inc., Natick MA, USA). These muscles were selected because, given the arm configuration used in this study and based on previous studies (De Comite et al., 2021), they are strongly recruited to counter rightward and leftward step loads.

**Figure 1:**
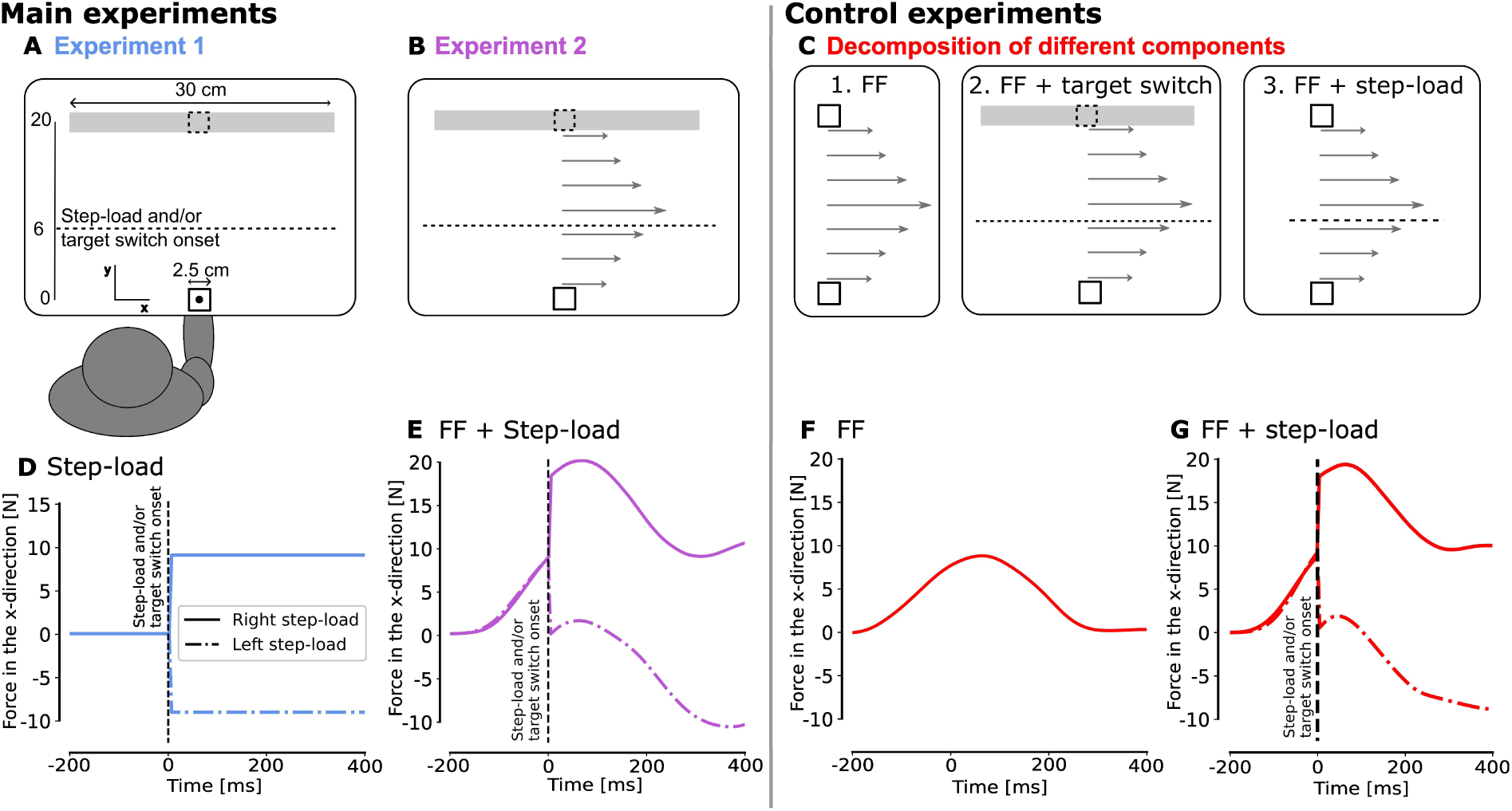
Experimental paradigm. **A**. Illustration of targets and forces for the first main experiment. Participants were asked to perform reaching movements of 20 cm between the home target and the goal target, which could either be a rectangle or a square. For trials with target switch, the change occurred during the movement (from rectangle to square or vice-versa). For trials with mechanical perturbations, rightward or leftward step loads were applied during the movement. The onset of the target switch and step loads is represented by the dotted line at 6 cm in the y-direction from the home target. This line was not visible to the participants, who saw only targets and a cursor representing the position of their hand. **B**. Schematic of the second main experiment. The instructions were the same as for the first experiment but in this case, participants were exposed to a velocity-dependent force field (𝐹_x_ = 13𝑦· [N]) throughout the experiment. **C**. Schematic of the control experiments. The different components of the second main experiment were decomposed in three control experiments: 1. FF only: participants were asked to perform reaching movements of 20 cm between the home target and the goal target, which was always a square, while exposed to the same velocity-dependent FF as in main Experiment 2, 2. FF + target switch: participants were again exposed to the same FF but, the goal target was a square or a square switching to a rectangle (randomly alternating), 3. FF + step-load: participants were again exposed to the same FF and performed their movement toward a square target but rightward or leftward step loads could be applied. **D-E-F-G.** Commanded force profiles during one exemplar trial for each experiment. Panels D, E and G represent the case of a right or left step load.

### Experimental design and statistical analysis

For all the experiments, each trial followed the same sequence. At the beginning of the trial, the home and goal targets were displayed in grey. Participants had to move the hand-aligned cursor to the home target, which turned green when participants reached it. The goal target became blue when participants were allowed to initiate their movement. This go-cue was triggered after a random delay uniformly distributed between 2s and 4s after reaching the home target. After this go-cue, participants could initiate their movement whenever they wanted so there were no constraints on the reaction time. The trial was successful if participants reached the goal target between 250 and 550ms after they left the home target and stabilised in it for at least 500ms. If the trial was successful, the goal target turned green, and the score projected onto the workspace was incremented by 1 point. If the movement was too slow or fast (movement duration outside the time window), the goal target turned red or black with a red outline, respectively. If the target was missed, it turned grey again. In these three last cases, participants did not score any points. After the feedback was given to the participants, all targets disappeared for 750ms and then reappeared for the next trial. Only successful trials were kept for kinematics analysis of Experiments 1 and 2. This represents 85.3% ± 7.49 (mean ± SD) of trials for Experiment 1 and 81.5% ± 10.76 (mean ± SD) of trials for Experiment 2. However, the results obtained are similar when all trials are considered. For analyses concerning the adaptation to the force field, all trials exposed to the force field were kept, even if they were too slow or fast.

#### Main experiment 1

This first experiment was an adaptation of an experiment performed by De Comite and colleagues (De Comite et al., 2021). In Experiment 1 (Figure 1A), participants were instructed to perform reaching movements from the home target (2.5 cm x 2.5 cm) to the goal target located at 20 cm in the y-direction that could be either a small square (2.5 cm x 2.5 cm) or a wide rectangle (30 cm x 2.5 cm). The wide axis of the goal target was orthogonal to the straight-line path from the home target to the goal target.

During one trial, participants could face two types of online perturbations, either separated or combined: a lateral step load applied by the robot on their hand, or a visual change in goal target shape, called target switch. The magnitude of the step load was +/-9N (rise time of 5ms), aligned with the x-axis. This force was triggered based on a hand position criterion, i.e., when the participant’s hand crossed a virtual line parallel to the x-axis and located 6cm from the starting point. It was switched off at the end of the movement (after the stabilisation period). The target switch was triggered at the same time as the step load and consisted of an instantaneous change of the target shape, from a square to a rectangle, or vice-versa.

The experiment was divided into 8 blocks of 58 trials. Each block included 18 trials without target switch or step load (9 trials per target type: square or rectangle), 16 trials with only a step load (4 trials per combination of target type (2) and step load direction: left or right), 8 trials with only a target switch (4 trials per switching target: square to rectangle and rectangle to square) and 16 trials with both a step load and a target switch (4 trials per target switch type (2) and per step load direction (2)). In total, participants performed 464 trials, including 32 trials for each combination of perturbations (except for the no-perturbation trials, where 72 trials were included). A score corresponding to the number of successful trials was projected on the virtual display to motivate participants. All these trials were randomly presented to the participants except for the first block, which began with 8 trials towards a square or a rectangle without target switch or step load, so the participants could start with easy trials. Before these 8 blocks, participants performed a training block composed of 24 trials, 2 of each type. In total, the experiment took two hours including explanation and preparation of the participant for the experiment.

#### Main experiment 2

The second experiment was very similar to the first one, with the addition of a velocity-dependent force field (FF) (Figure 1B). This velocity-dependent force field was applied to the hand of the participants in addition to the other perturbations. This force field consisted of a lateral force to the right proportional to the forward hand velocity: 𝐹_x_ = 13𝑦·. We chose to use a single-component field for simplicity, as the direction of adaptation then aligns with the rectangle in the bar task under the assumption that adaptation patterns to orthogonal and curl fields are similar (Crevecoeur et al., 2020a; 2020b). Some catch trials in which the velocity-dependent force field was removed were inserted randomly in about 9% of trials (48 trials on 512).

In this experiment, participants performed 8 blocks of 64 movements. The distribution of trials was the same as in Experiment 1, with 6 additional catch trials per block, 3 towards a square and 3 towards a rectangle. Before the experiment, participants were exposed to a training block without FF, as for the first experiment. The first block started with 8 trials towards a square or a rectangle without other perturbations than the velocity-dependent force field. This was done to avoid the first trials with step-load and/or target switch being missed. The experiment also took two hours in total.

#### Control experiments

Three control experiments were designed to decompose all the different factors present in Experiment 2. This first control experiment was a classical adaptation experiment in which participants had to perform reaching movements while exposed to a velocity-dependent force field similar to the one used in Experiment 2 (Figure 1C1). Again, participants started from the home target (2.5 cm x 2.5 cm) and aimed for a similar goal target presented as a square at the same location and identical to the one used in Experiments 1 and 2 (2.5 cm x 2.5 cm, 20 cm in the y-direction). The second control experiment (Figure 1C2) was the same as the previous one with the addition of target redundancy with or without a switch. Indeed, 6 participants experimented with a rectangle goal target in half of the trials and six participants experimented with the goal target which was a square switching to a rectangle in half of the trials (the other half being a square target). As the results in terms of motor adaptation for these two experiments were similar, the two experiments were merged. The last control experiment was also a classical adaptation experiment with the addition of a step load to the left or the right. These step loads were identical and in the same proportion as in Experiment 2 (Figure 1C3).

In the three control experiments, participants started with a first block of 30 trials without force field to get used to the task. Then, they performed 5 blocks of 50 trials in the presence of the force field (and the corresponding additional perturbations). Each of these blocks included 5 catch trials, randomly inserted (10% of trials). Because we were interested in feedback corrections, we did not use error clamps or force channels, as these types of trials constrain movement and suppress natural corrective responses, limiting our ability to assess how participants responded to perturbations. At the end of the experiment, participants performed a wash-out block of 30 trials without force field. The experiment took one hour in total.

With the main and the control experiments together, we can analyse, on the one hand, the influence of motor adaptation on policy update by comparing the results of the main experiments, and, on the other hand, the influence of target switch and step loads on motor adaptation by comparing Experiment 2 with the data from the different control experiments.

#### Data Analysis

Kinematics data were recorded with a sampling rate of 1kHz using Kinarm’s Dexterit-E software (Version 3.9). Preprocessing of the data was done using custom MATLAB scripts (MATLAB 2022a, Mathworks Inc. Natick Ma, USA). Kinematics data were filtered with a dual pass 3^rd^ order low-pass Butterworth filter with a cutoff frequency of 20 Hz. The rest of the analyses were done using Python 3.11 via Anaconda Software Distribution (Version 23.7.4; Anaconda Inc. 2023). All kinematic data were aligned with the onset of the step load (which also corresponds to a position threshold, as the step load is always applied when the hand crosses a virtual horizontal line located at 6cm from the home target). We defined the final hand position as the position of the hand when the velocity dropped below a threshold set to 0.05𝑣_max_ during 150 ms, where 𝑣_max_ was the maximal velocity observed during the corresponding movement.

For the main experiments, the activity of two muscles of interest was measured. The two chosen muscles were the Pectoralis Major (PM) and the Posterior Deltoid (PD) in the right shoulder, as these are the main muscles recruited to respond to the left and right step loads in this configuration. Before placing the electrode over the muscle belly, the skin of the participants was cleaned with cotton wool and disinfected with medical alcohol. Gel was applied to the electrodes to improve signal conduction.

Muscular data were sampled at 1kHz and amplified with a factor 10^4^. The first preprocessing step for muscular data was done using custom MATLAB scripts (MATLAB 2022a, Mathworks Inc. Natick Ma, USA). Data were filtered with a 6th-order Butterworth band-pass filter with cutoff frequencies [20,250] Hz, then exported to be analysed using Python as for the kinematic data. Muscle recordings were normalised to the average activity of each muscle sample during three separate calibration blocks done at the beginning of the experiment and after the third and sixth blocks. During these calibration blocks, a 2.5 by 2.5 cm square was presented to the participants. They had to keep the cursor within this target. As soon as they entered it, a right or left step load of 9N was applied to their hand to activate one of the muscles of interest. Participants were asked to counter the force to come back to the target. The force was applied during 2.5s and 8 times in each direction. For each step load of the block, we took the muscular activity between 0.5s and 2s following the onset of force to compute the mean muscular activity during the calibration. The muscular activity during the whole experiment was divided by the calibration value corresponding to the muscle and aligned on the force onset.

When analysing the data, we primarily focused on trials where participants experienced a step load to the right, in the same direction as the FF. This approach was taken because, when participants were exposed to a leftward step load, as the FF is directed to the right, the effect of the step load was attenuated, thereby reducing the statistical power of the analysis. However, these leftward perturbed trials played a critical role in preventing participants from anticipating the step load direction before movement. Moreover, for the kinematics, we selected the subsets of data that most directly addressed our central hypothesis and focused on trials with a target switch from a square to rectangle. This allowed evaluating the ability of participants to take advantage of the target redundancy.

#### Statistical analysis

The statistical difference between groups across Experiments 1 and 2 (with or without FF) for the final hand position and the difference in variance was computed using student’s t-tests. The difference was considered significant when *p < 0.05* as we made an inter-subjects comparison and used EMG data.

To evaluate feedback responses depending on the change in target shape (square vs square to rectangle target) in both groups, we analysed normalised EMG data. We only looked at the stretch response in the agonist muscle to the direction of the step load. The EMG data of each participant were averaged across trials of the same condition. Data were then averaged with non-overlapping bins of 25 ms width. This bin width was the same as the one usually selected in other similar studies (Kurtzer et al., 2008; Nashed et al., 2012; De Comite et al., 2021).

A linear mixed model was used to assess the statistical difference in muscle activation and in final position. The general equation of this linear mixed model is :

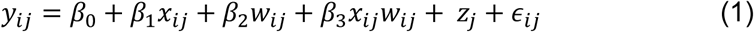

where 𝑥_ij_ 𝑎𝑛𝑑 𝑤_ij_ were the fixed effect factors and 𝑧_j_ the random effect factor (participants). To assess the difference in final position depending on the number of trials performed (𝑥_ij_) and the group of participants (𝑤_ij_), we focused on the coefficients of these fixed effect factors, namely 𝛽_1_and 𝛽_2_ respectively, as well as the interaction term 𝛽_3_, which captures whether the effect of the number of trials differs between groups. To assess the difference in muscular activity between trial types (𝑥_ij_), we looked at the fixed effect factor 𝛽_1_. In that case, we made comparisons between participants in the same group, and have therefore disregarded the group effect (𝑤_ij_). In both cases, the difference was considered significant when *p < 0.05*.

For the characterization of motor adaptation when the experiments involved a force field, we extracted the maximal deviation, which is the most eccentric position in the direction of the force field. This maximal deviation usually decreases with exposure to the FF and can be fitted with an exponential regression. A bootstrapping technique was used to resample participants (with replacement) one million times. The exponential regression model:

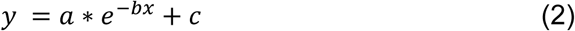

was fitted to each resampled dataset to obtain distributions for the parameters 𝑏 and 𝑐. This procedure allows for the estimation of the variability of the regression parameters across the different resamples, providing confidence intervals and assessing the stability of the model’s estimates. Because the obtained distributions for b and c parameters were non-normal, the comparisons between groups were based on their 95% Highest Density Intervals (HDI) (Murphy, 2012).

## Results

### Experiments 1 and 2: influence of the FF on flexible control

We first compared the two main experiments to assess whether exposure to the FF affected participants’ ability to react to online changes (visual or mechanical). In the first experiment, participants were exposed to changes in target shape during movement and step loads applied either leftward or rightward. The second experiment mirrored the first one but included a clockwise velocity-dependent FF. Figures 1D and E show the force profile applied to the hand during movement in the case of a right or left step load for Experiments 1 and 2, respectively. Figures 2A, B, F, and G depict the kinematics traces corresponding to rectangle and square goal targets without any target switch. As previously reported (De Comite et al., 2021), participants’ behaviour in Experiment 1 depended on the shape of the target. A significant difference in intra-subject variance was observed between the square target (endpoint variance = 0.127) and the rectangle target (endpoint variance = 1.704, t(19) = 3.278, p = 0.004, paired t-test). This larger variance for the rectangle target was indicative of the participant’s tendency to exploit target redundancy.

**Figure 2:**
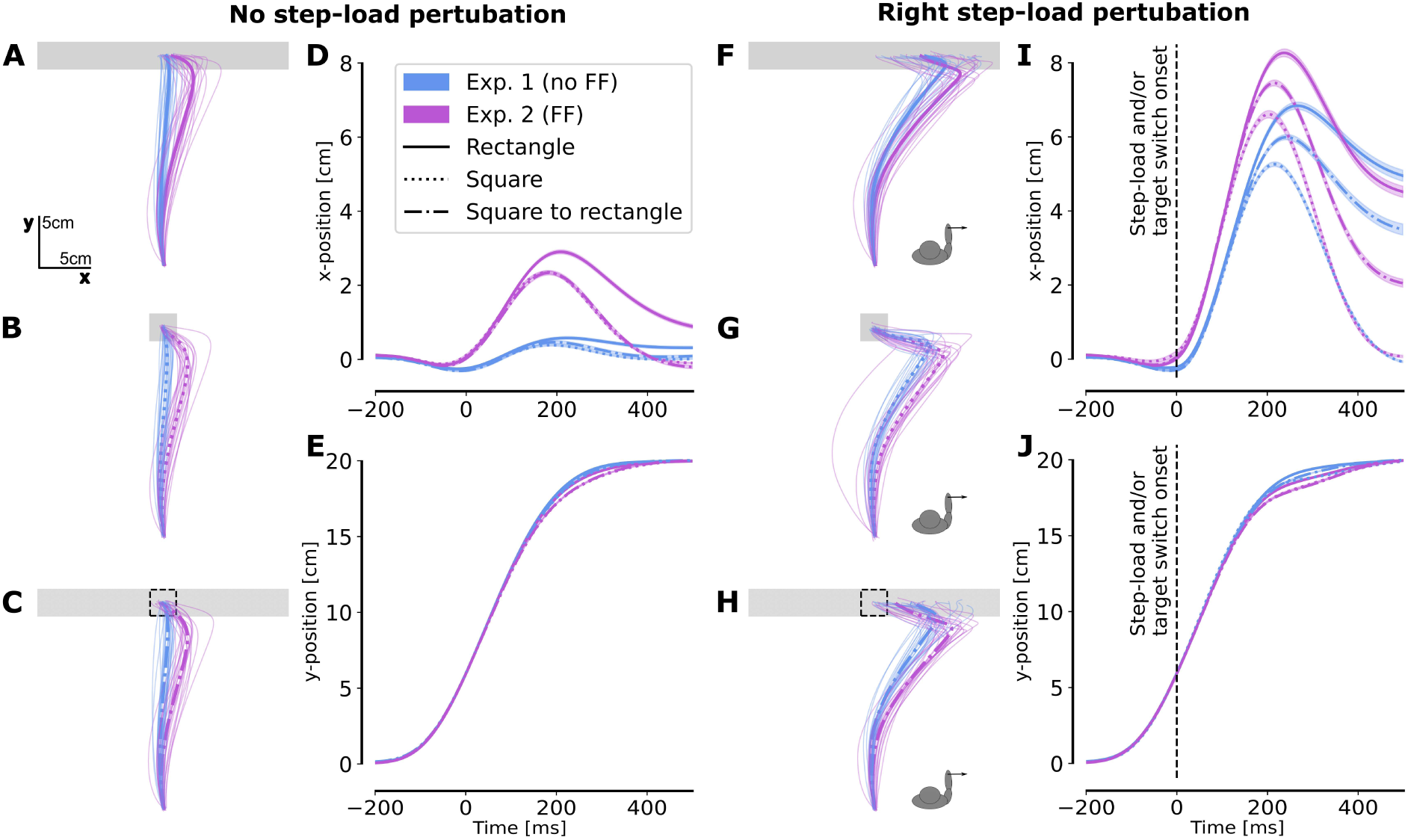
Kinematics. **A-B-C.** Mean x-y traces across trials for each participant (light traces), and across participants (thick trace) with the FF (Exp. 2, purple), or without FF (Exp. 1, blue) from trials in which there was no step load for **A.** a rectangle goal target, **B.** a square goal target, **C.** a square goal target initially which became a rectangle during movement. **D.** Evolution of the x-position (mean across participants ± SE) over time during the movement with FF (purple), or without FF (blue) for a rectangle target (solid line), a square target (dotted line) or a switch from a square to a rectangle target (dashed line). Note that the curves for the square target and the switch from a square to a rectangle target are nearly superimposing each other. **E.** Evolution of the y-position (mean across participants ± SE) over time during the movement for all conditions. Note that all the curves are nearly superimposing each other **F-G-H-I-J.** Same as panels A to E for trials with a right step load.

Similar observations were made regarding the endpoint variance when participants performed movements in the presence of the FF, with the intra-subject variance remaining significantly different (endpoint variance for the square = 0.123, endpoint variance for the rectangle = 2.983, t(19) = 4.999, p = 7.97 x 10^−5^, paired t-test). The difference in endpoint variance was even more pronounced in this case as the FF deviated the participants’ hand to the right. The x-position during movement increased more (Figure 2D) and participants exploited the target’s redundancy by letting their hand deviate along the target axis as indicated by the increase in variance in this direction. Together, these increases in final x-position and variance indicate a motor strategy that exploits the structure of the task. A more detailed analysis of the final position and the evolution of variance is presented later (see Figure 3).

**Figure 3:**
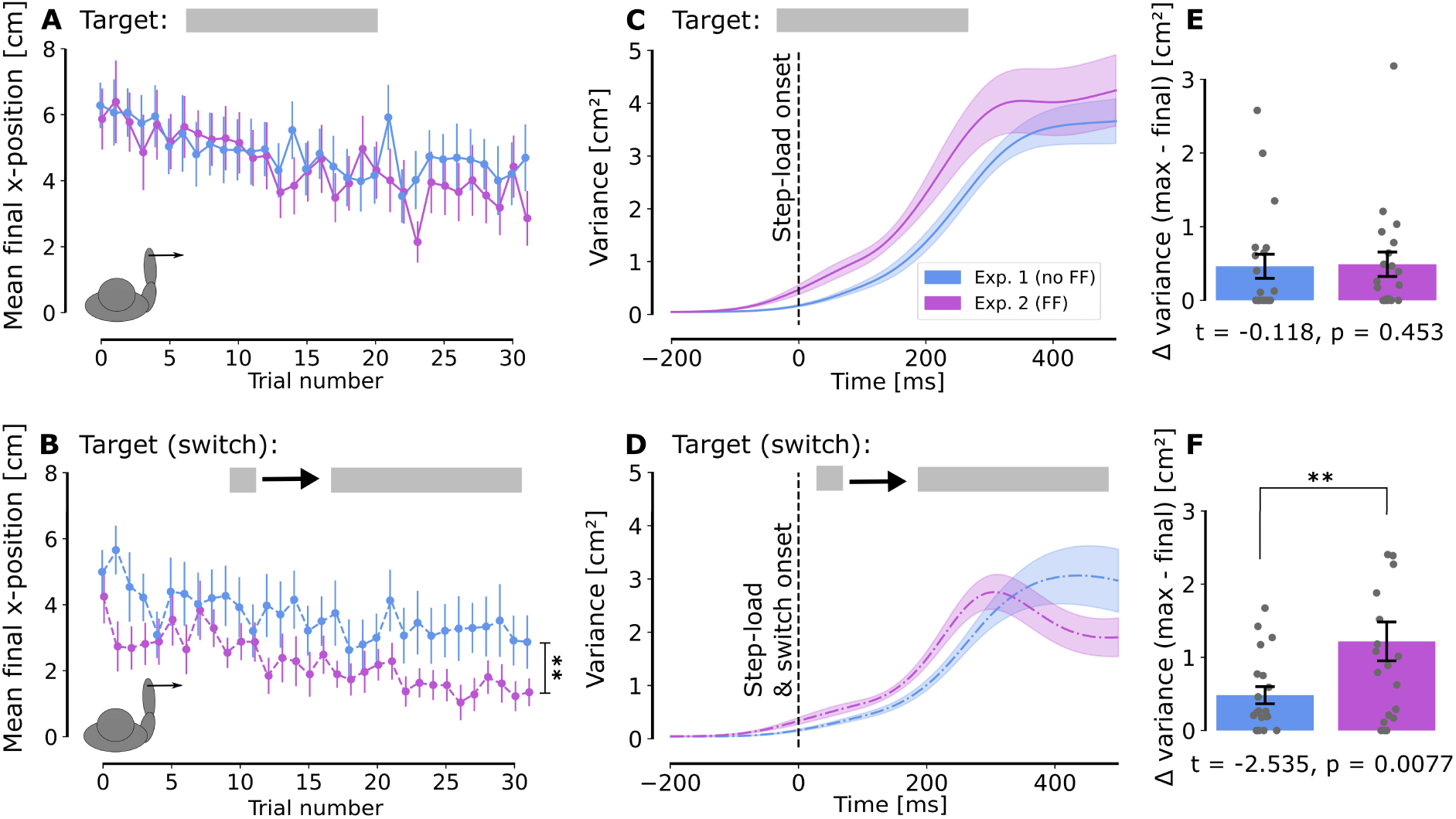
Exploitation of target redundancy. **A-B.** Average final x-position along the rectangle width without force field (blue) and with force field (purple) following a right step load for each trial (mean across participants ± SE) when **A.** the goal target was a rectangle and there was no visual switch, **B.** the goal target switched from a square to a rectangle. **C-D.** Evolution of intrasubject variance with time (mean across participants ± SE) when **C.** there was no visual switch of the rectangle target, **D.** the goal target switched from a square to a rectangle. **E-F.** Mean of the difference between the maximal and the final variance for each experimental group when there is **E.** no switch of the goal target, **F.** a switch from square to rectangle. Positive values mean the endpoint variance was smaller than the peak variance across trials.

We can then look at the movements in which participants are exposed to rightward or leftward step load with or without FF. The endpoint variance was again higher for the rectangle target than for the square target for all conditions of step loads being with or without FF. For left step load without force field (endpoint variance square = 0.141, rectangle = 5.58; t = 6.826, p < 4.657 x 10^−10^, paired t-test) and with force field (endpoint variance square = 0.104, rectangle = 4.74; t = 6.646, p < 2.34 x 10^−6^, paired t-test). For right step load, without force-field (endpoint variance square = 0.146, rectangle = 5.653; t(19) = 6.486, p < 4.23 x 10^−6^, paired t-test) and with force-field (endpoint variance square = 0.112, rectangle = 6.82; t(19) = 7.791, p = 2.48 x 10^−7^, paired t-test). These findings collectively indicate that when the target was a rectangle, participants took advantage of target redundancy, resulting in higher endpoint variance.

No differences were observed in the y-position across conditions (Figure 2 E, J). However, regarding the x-position when participants were exposed to a rightward step load (Figure 2I), the deviation in the positive x-direction was higher at the beginning of the movement with the FF. However, the final x-position was higher without FF, regardless of whether the target is a rectangle or a square transforming into a rectangle during movement. This is indicative of a bigger correction towards the centre of the rectangle target in the presence of the FF.

We can then focus on the final position in the target. Under a step load without a target switch, participants effectively exploited target redundancy. Specifically, under rightward step loads, participants utilised target redundancy similarly, irrespective of the presence of the force field. This observation serves as a control condition: even if the net total force was different in Experiment 2 as the resultant was the sum of the force field and step-load, our data indicated that participants exploited target redundancy in the same way when there was no target switch. Based on this condition, the effect of the switch could be interpreted in spite of different total forces across experiments. Figure 3A shows the evolution of the final position across all trials in this condition (rectangle target, no switch, right step load). A reduction in endpoint eccentricity across trials was observed. A linear mixed model was fitted to further characterise the evolution of the final position used as a dependent variable as a function of the trial number and group as fixed predictors and a random offset for each participant (see Methods). Only the final positions of valid trials were considered. For the trials towards a rectangle target without a switch, the effect of the trial number was significant (𝛽_1_ = -0.05, CI 95% [-0.082, -0.018], z(1131) = -3.091, p = 0.002), with no significant effects observed for the group (𝛽_2_ = 0.233, CI 95% [-0.593, 1.060], z(1131) = 0.554, p = 0.580) or the interaction between groups and trial number (𝛽_3_ = -0.038, CI 95% [-0.083, 0.007], z(1131) = -1.664, p = 0.096). However, when a switch from a square to a rectangle target occurred (Figure 3B), both the trial number (𝛽_1_ = - 0.055, CI 95% [-0.08, -0.03], z(1121) = -4.362, p = 1.29 x 10^−5^) and the group (𝛽_2_ = -1.019, CI 95% [-1.667, -0.37], z(1121) = -3.079, p = 0.002) had significant effects, with no significant interaction (𝛽_3_ = -0.022, CI 95% [-0.057, 0.014], z(1121) = -1.198, p = 0.231). Thus, the final position on the target was significantly less eccentric when participants encountered both the FF and a visual switch in the target shape. Interestingly, this difference in the final position on the target was not observed without a target switch, suggesting a reduced ability of participants to take advantage of target redundancy because of the combination of sudden change in target shape and exposition to the FF and not due to the FF alone.

A reduced ability to exploit target redundancy was also highlighted by the temporal profile of the variance along the x-axis. Indeed, a reduction in variance reflects the use of a state-dependent control policy that steers the hand back towards a target state (Todorov & Jordan, 2002b). Interestingly, such a decrease in intra-subject variance was observed when participants encountered the combination of FF and target switch (Figure 3D) but not when there was no switch in target (Figure 3C). We extracted the difference between the maximum variance across trials for each participant and the endpoint variance in this scenario. A positive value indicates that the variance peaks during movement and decreases near the endpoint, whereas a value of 0 indicates that the peak variance is observed at the endpoint. We observed that the endpoint variance was significantly reduced when participants were exposed to the FF in the switch condition (t(38) = 2.535, p = 0.0077, one-tailed independent t-test) (Figure 3F), whereas no significant difference was observed without the target switch (t(38) = -0.118, p = 0.453, one-tailed independent t-test) (Figure 3E). This further illustrates participants’ reduced ability to exploit target redundancy as the reduction in variance when exposed to FF and target switches indicated a tighter control of lateral coordinates in spite of the target redundancy.

Muscle activities related to step loads were analysed by comparing mean activity during perturbed trials to that during unperturbed trials. Consistent with previous reports (De Comite et al., 2021), muscle activity towards a rectangle target was lower compared to a square target. In this study, the contrast of muscular activity between the movements toward a square or a square switching to a rectangle for Experiments 1 and 2 (Figure 4 A and D) was analysed. In this analysis, the muscular activity in response to the FF has been removed when subtracting the muscular activity during trials without step loads. However, the raw muscular activity before step load is higher when participants are exposed to the FF (mean in a 50ms window before onset of step load: 0.54 [a.u] without FF, 0.98 [a.u] with FF). This leads to gain scaling in short-latency stretch responses as the same muscle stretch will elicit larger responses for larger preperturbation muscle activity (Pruszynski et al., 2009). For Experiment 1, a clear difference in muscular activity emerged in the voluntary window (100-180ms after force onset), which was less pronounced in the presence of the FF. The data were then grouped into 25 ms bins, aggregating the data over non-overlapping 25 ms time windows. Using a linear mixed model on mean muscular activity across these 25 ms bins (Figure 4 B, E), a statistically significant difference was found from 100ms to 200ms (p < 0.005 for all bins). This difference was still present, although less significant; for some bins after 100 ms when the force field was present. By looking at the evolution of p-values over a 25ms sliding window, a noticeable drop could be seen after 100 ms without FF in Experiment 1 (Figure 4C), whereas it was less clear with FF (Figure 4F). The reduced statistical significance observed in the FF condition is due to a meaningful attenuation of the effect—consistent with our behavioural findings. Specifically, the movement kinematics showed that participants were still able to exploit target redundancy when they were exposed to the FF, only to a lesser extent. The EMG results mirrored this pattern. To avoid the arbitrary binning and consider correction for multiple comparisons, a permutation test was performed in addition. For both Experiment 1 and 2, within-subject difference was computed between the two conditions (square and switch from square to rectangle) at each time point. For each permutation iteration, the sign of the difference was randomly flipped for each participant, simulating the null hypothesis of no difference across subjects. Then, the mean difference across subjects was computed for each time point, building a null distribution of differences over 1000 permutations. For each time point, a two-sided p-value was computed by comparing the real mean difference to the null distribution and the p-value was corrected for multiple comparisons using the False Discovery Rate correction. The results of this analysis indicated a cluster of significant differences starting a bit before 150ms without FF and no significant difference with FF. These results confirm our previous observation, showing a weaker effect of the observed difference when participants were exposed to the FF. Figure 4G illustrates distinct differences in muscular activity depending on the goal target during the long-latency and voluntary windows. In both experiments for the voluntary window, a gradual decrease in muscle activity was observed from the square target to the rectangle target. The muscular activity for a square target and a rectangle becoming a square was significantly higher than for a square becoming a rectangle or a rectangle, being without or with FF (even if it is less significant with FF). Interestingly, this clear modulation of the muscular activity depending on the target showed that participants were able to flexibly adjust their motor response. This indicates that there was no systematic up-regulation of co-contraction used as a strategy to enhance limb impedance and robustness irrespective of the target geometry (Burdet et al., 2001; Crevecoeur et al., 2019). We further checked that participants exposed to the FF did not exhibit muscle co-contraction before step load onset. A significant difference was observed in pectoralis activity (t(38) = -2.559, p = 0.015, two-tailed independent t-test) indicating a response to the velocity-dependent FF, already active before step load onset. However, no significant results were found for the deltoid activation (t(38) = - 0.286, t = 0.777, two-tailed independent t-test), suggesting no additional muscle contraction as the default strategy against perturbations.

**Figure 4:**
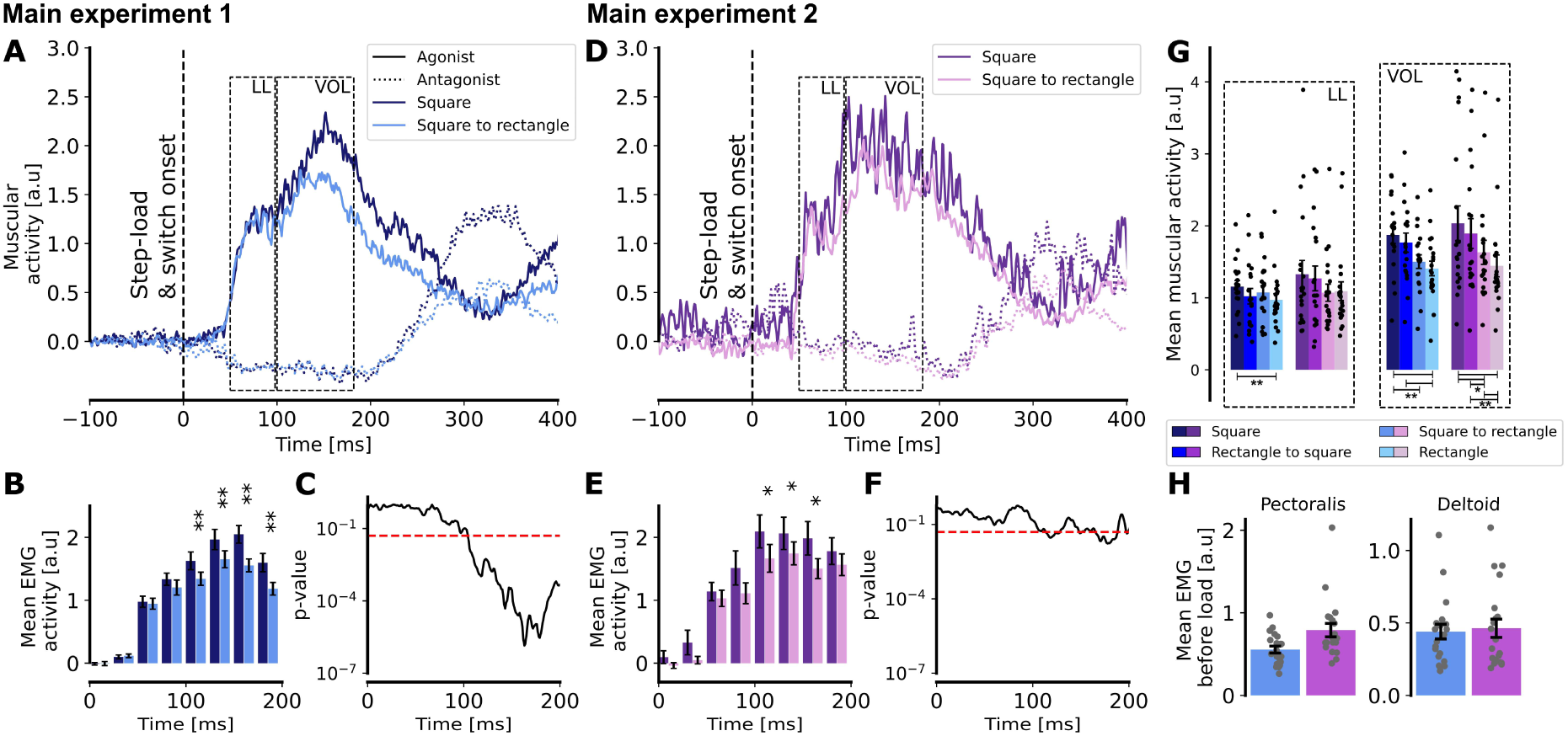
Muscular activity. For the first main experiment (without FF), **A.** muscular activity in the Pectoralis Major (solid line) and in the Posterior Deltoid (dotted line) following a right step load when the target was a square (dark blue) or a square which became a rectangle (light blue). **B.** Mean muscular activity over 25ms bins for a square target (dark blue) or a target switching from a square to a rectangle (light blue). **C.** Log-scale evolution of the p-value over a 25ms sliding window. Time index corresponds to the timing of the end of the window. Red horizontal line represents p = 0.05. **D-E-F.** Same as A-B-C, but for the second main experiment, with FF. **G.** Mean muscular activity for each target condition during the long latency and the voluntary windows. **H.** Mean muscular activity across all trials in the Pectoralis Major and Posterior Deltoid in a window of 300ms before the step load for the experiment without FF (blue) and with FF (purple).

These findings collectively indicate that participants were able to exploit target redundancy while adapting to the FF, but that this ability was reduced when participants were exposed to the FF in addition to a target switch. The final position on the target was significantly less eccentric and a decrease in variance at the end of the movement was observed. These results were reproduced by looking at the muscular activity. The absence of systematic co-contraction suggests a modulation of control gains during movement at latencies that were a bit shorter than the ones previously reported (De Comite et al., 2021). These results therefore highlight a specific interference between flexible control (represented here by the target switch during movement) and motor adaptation.

### Experiment 2 and control experiments: Reduced extent of adaptation

Next, we sought to analyse the potential effect of changes in target redundancy on motor adaptation. To explore this, we compared the data from the second main experiment, where participants encountered online control and a velocity-dependent FF, to the data from the first control experiment featuring only the FF. The data from the second experiment were sub-sampled to extract only the trials towards a square target and analyse them to their actual trial indices. This allows comparing trials to the same target by taking into account the number of trials participants were exposed to the FF previously. The time axis was thus not influenced by this sub-sampling and so, a similar learning rate would indicate that the intermediate trials with other perturbations (of target or forces) did not slow down the adaptation. Figure 5A shows the learning curves from these experiments. Notably, the experiment incorporating both switches in target shape and step loads had twice the number of trials. It can be observed that participants reached a lower maximal deviation during standard FF adaptation. The after-effect quantified by the deviation in the direction opposite to the FF during catch trials was significantly more pronounced with the FF alone (t(30) = 3.868, p = 5.47 x 10^−4^, two-tailed independent t-test, Figure 5B). To quantify differences in learning rate and extent of adaptation, an analysis of the 𝑏 and 𝑐 parameters of the exponential regression (see Methods) was conducted (Figure 5C). The bootstrap analysis revealed an overlap in the 95% HDI for the 𝑏 parameter, indicating a similar learning rate across both experiments. However, there was no overlap in the 95% HDI for the 𝑐 parameter indicating a reduced extent of adaptation in Experiment 2. The parameter *a* was not shown as it is directly linked to the *c* parameter. Indeed, the sum *a+c* corresponds to the starting point of the exponential curve, which was similar in both cases as evidenced by the overlap of 95% HDI for the distribution of *a+c*: [3.7, 4.94] in the FF condition and [3.35, 4.52] in the FF + step load + switch condition.

**Figure 5:**
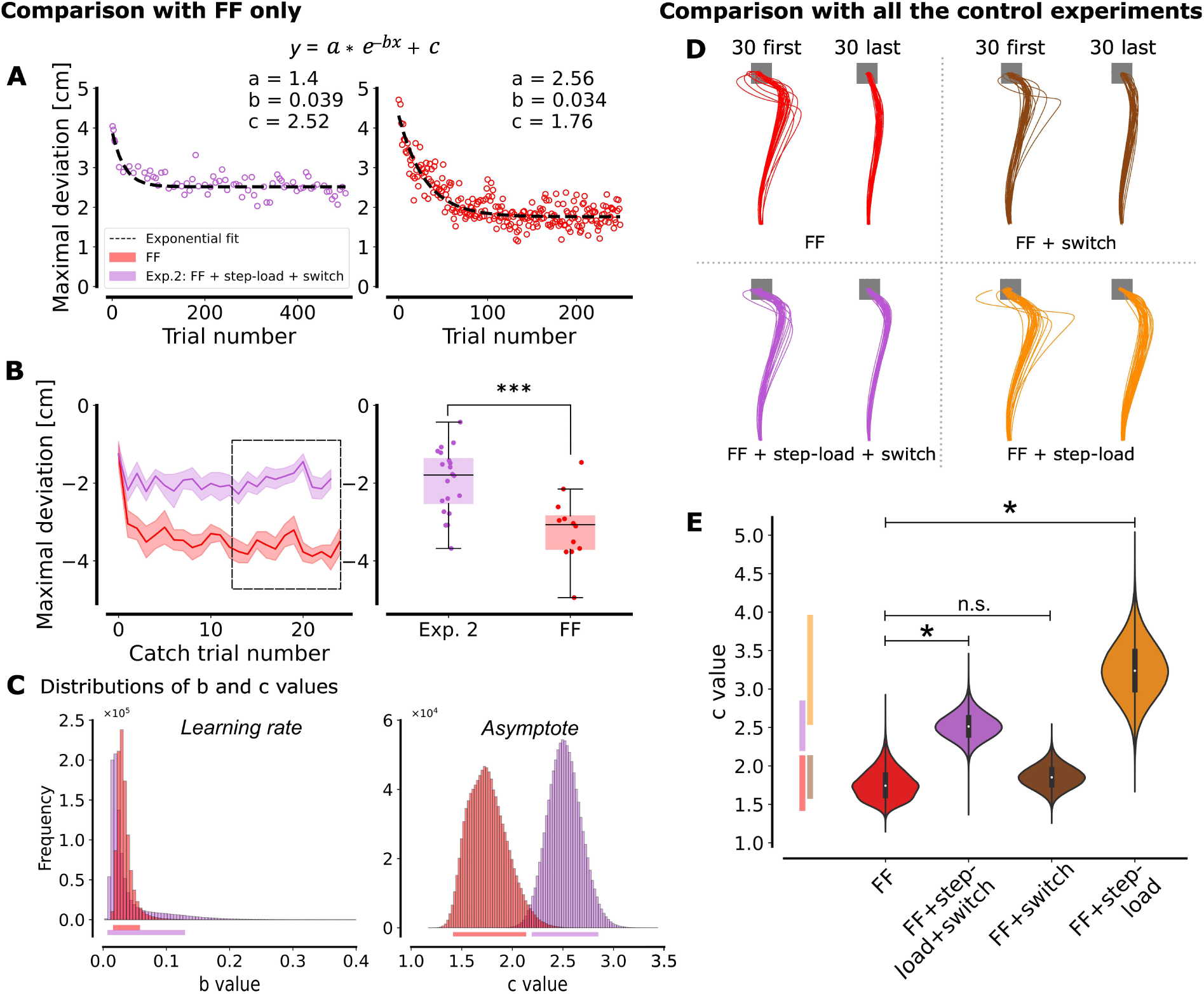
Motor adaptation. **A.** Maximal deviation (with respect to the straight line between the home and goal target) across trials for which participants are exposed to the force field for the experiment with target switch and step load in addition to the force field (purple) and the control experiment with the force field only (red). Trials towards a square target are represented against their index. **B.** Maximal deviation during catch trials represented against their index for both experiments (mean across participants + SEM) (left part) and median deviation across the ten last catch trials (right part). **C.** Distribution of the 𝑏 and 𝑐 parameters of the exponential regression after 10^6^ resampling of the population with replacement. Horizontal lines below the graphs represent the 95% highest density intervals. **D.** Comparison across all control experiments. Mean traces for the first (left) and last (right) 30 trials exposed to the force field for the experiment with the force field only (red), the experiment with target switch and step loads in addition to the force field (purple), the experiment with only target redundancy (rectangle and switch from square to rectangle, brown) and the experiment with step loads in addition to the force field (orange). Each line represents the mean for one trial across participants. **E.** Value of the 𝑐 parameter of the exponential regression for the same four experiments. The 95% highest density intervals are projected on the y-axis.

Further experiments were warranted to elucidate whether this reduced adaptation stemmed from the target shape or the step load. To explore this further, two additional control experiments were conducted to separate the effects of the target shape and the step load. Figure 5D illustrates the mean trajectories for the first 30 and last 30 trials from Experiment 2 and from each control experiment. Initial trials exhibited substantial deviation, but varying levels of residual deviation towards the FF direction were observed in the 30 last trials across conditions. This was reflected by the 𝑐 value of the exponential regression (Figure 5E). The HDI overlap between the FF and FF + switch experiments indicated that the residual deviation, and therefore, the extent of adaptation was similar in these two conditions. This indicated that changes in target shape did not influence the extent of adaptation to the FF. A dedicated analysis of behaviour during learning with a rectangle target would require a follow-up study. However, an overlap of HDI for the FF + step load experiment with the corresponding interval in purple (Exp. 2) was observed. This overlap indicated that when we added a step load, the extent of adaptation was reduced.

Altogether, these results indicate that the learning rate is similar between the standard adaptation to the FF and the second experiment in which we added target switches and step loads. A difference in the extent of adaptation was however observed between these two experiments and further control experiments indicated that this difference in the extent of adaptation stemmed from the step loads.

We thus demonstrated an asymmetric interference between motor adaptation and flexible control (considered here as adjustments to online changes, i.e., switch in target or step loads). On the one hand, motor adaptation influenced flexible control by reducing participants’ ability to exploit target redundancy when the target switched from a square to a rectangle. On the other hand, flexible control (and more specifically step loads) reduced the extent of adaptation to the FF.

## Discussion

This study investigated the interplay between flexible control and motor adaptation to determine whether these processes were completely independent, or whether any interdependence between them could indicate possible overlap in their neural bases. We compared participants’ behaviour in tasks involving either flexible control, motor adaptation, or both. Notably, we observed distinct behaviours when both mechanisms were engaged simultaneously compared to when they were engaged separately. We made the following key observations: first, when both mechanisms were combined, participants exhibited a reduced ability to exploit the redundancy of the rectangle target; second, they also demonstrated a lesser extent of adaptation to the force field.

The results from the first experiment were consistent with previous studies on the influence of target redundancy on reaching control policies (Nashed et al., 2012; De Comite et al., 2021). The difference in endpoint variance depending on the target shape indicated a modulation of the feedback gains depending on target redundancy. This modulation was further reflected in the EMG data, which showed a modulation consistent with the target structure and provided important measurements about the timing of this modulation. The latencies observed for EMG responses in Experiments 1 and 2, around 100ms, are comparable to the latencies of the visuomotor system (S. Franklin et al., 2017) and align with the findings that response amplitude to target jumps is sensitive to goal redundancy within ∼90 ms (Cross et al., 2019). Unexpectedly, these latencies were shorter than the ∼150ms reported by De Comite et al. (2021), suggesting that the time needed to change the control policy could have been overestimated in this previous study potentially due to a smaller sample size. The analysis of participants’ ability to exploit target redundancy across trials revealed a progressive reduction in lateral endpoint eccentricity, regardless of the presence of a FF (Figures 3A, B). This aligned with Orban de Xivry and Lefèvre (2016), who found that random target width changes reduced exploitation of target redundancy, possibly reflecting a switching cost. The effect was similar across conditions with or without FF, as reflected by the comparable slopes for lateral eccentricity evolution (Figures 3A, B).

Regarding motor adaptation (Figure 5, red curves), our results aligned with previous studies, showing an exponential decay of the maximal deviation in the FF direction (Shadmehr and Mussa-Ivaldi, 1994; Dizio and Lackner, 1995; Lackner and DiZio, 2005; Cluff and Scott, 2013). This observation was based on a statistical fit of a standard exponential model, in which we did not make any assumptions about the underlying dynamics. In contrast, other approaches were previously used in the literature. For example, state-space models (Smith et al., 2006; Kording et al., 2007) describe learning as a balance between error-dependent correction and baseline reversion, a sort of forgetting (Vaswani et al., 2015). Besides, the Kalman filter model integrates optimally sensory feedback and state estimates (Wei and Körding, 2010). While our approach focused on a statistical description, incorporating these dynamics-based models in future work could clarify the contributions of error correction, sensory integration, and baseline reversion, providing complementary perspectives on motor adaptation.

The main contribution of our work is to highlight a specific interference in two identified conditions. Although the experimental paradigm resembles a dual-task setup, the interference observed was not trivial and was specifically linked to motor control processes. Our first main observation was that the presence of a force field limited participants’ ability to modify their controller following target switches. This observation provides indirect evidence that sensorimotor adaptation operates online as already suggested in previous studies (Crevecoeur et al., 2020a, 2020b). The first main experiment showed that target redundancy exploitation was consistent across conditions with or without FF when the target did not change (Figure 3 A, C). Under the hypothesis that sensory prediction errors update an internal model of movement dynamics between trials (Shadmehr and Mussa-Ivaldi, 1994; Shadmehr et al., 2010), it is tempting to conclude that flexible feedback control and adaptation are unrelated. However, such an interpretation would miss the fact that interference became clearly apparent when the target switched during movement. This indicated that the interference mechanism occurred when the target switched, suggesting a specific timing for these perturbations to produce interference, that is when they both happened during an ongoing movement. The observations are consistent with our hypothesis that adaptation also engaged online processing (Kalidindi & Crevecoeur, 2023). The fact that interference occurred at a specific timing underscores the importance of measuring the timescales of these processes.

Our second key finding was the reduced extent of motor adaptation observed in Experiment 2. Our control experiment (Figure 5D, orange curve) revealed that it was due to the superposition of different force perturbations (velocity-dependent FF and step load).

Conceptually, these observations are consistent with the findings of Singh and Scott (2003) who showed that under uncertainty, participants relied on “local” sensorimotor associations, mapping loads sensed at one joint to control updates at the same joint. For instance, participants misattributed elbow loads as linked to elbow velocity, even when they were proportional to the shoulder. It was necessary to de-correlate the two signals with a different exposure protocol to enable proper association between shoulder velocity and elbow loads. Similarly, in our experiment, step loads added to the FF may have disrupted FF identification, suggesting interference about contextual parameters as generalised with the contextual inference model (COIN) (Heald et al., 2021).

Our results can be interpreted within the framework of stochastic optimal control (SOC) initially introduced by Todorov and Jordan (2002b) to explain how humans achieve reliable success despite variations in motor behaviour. In this model, changes in the target shape correspond to modifications of the cost function, while motor adaptation updates the state-space representation matrices. Mathematically, these two mechanisms therefore impact different sets of parameters (Kalidindi and Crevecoeur, 2023). However, our findings suggest interference between these mechanisms in practice. Specifically, when participants were exposed to the FF, they tended to apply a larger correction to the centre of the rectangle target. This likely reflects an increase in feedback gains evoked when exposed to novel dynamical contexts (S. Franklin et al., 2017; Crevecoeur et al., 2019; Coltman & Gribble, 2020). Within the SOC framework, higher feedback gains imply a stronger correction in response to step loads. However, when a target switch was introduced in addition to the FF, the corrections became even larger. This suggests an interaction between the mechanisms governing the modulation of feedback gains and those involved in updating the controller following changes in the cost-function. We could therefore hypothesise that a general cost function governs the update of these mechanisms, ensuring limited parameter adjustments for global efficiency. Alternatively, a hierarchical control structure may prioritize certain parameters over others, dynamically allocating resources based on task demands. Future modelling work should account for these shared resources and potential priority rules to better capture the interplay between those mechanisms.

Building on this understanding, our results allow formulating the hypothesis that the neural structures involved in flexible control and motor adaptation are not independent, raising questions about which neural structures might mediate this interaction. One candidate locus of interference is the cerebellum, which is known to contribute to long-latency reflexes in response to unexpected mechanical perturbations (Strick 1983; Kurtzer et al., 2013) and has been associated with sensorimotor adaptation (Shadmehr and Krakauer, 2008; Spampinato et al., 2017; Carey, 2024). Considering that afferent volleys of somatosensory feedback about the presence of both the force field and the step loads converge to the same cerebellar structure, signal superposition could disrupt how errors are mapped onto command updates.

Besides, the basal ganglia are often hypothesised to be involved in the representation of costs (Shadmehr and Krakauer, 2008); potentially influencing movement selection and change in controller depending on target shape. Bostan and Strick (2018) showed that basal ganglia and cerebellum form a densely interconnected network, with cerebellar motor output affecting basal ganglia input via the intra-laminar thalamic nuclei. The reduced exploitation of target redundancy observed in our study during FF exposure may reflect a control policy shift driven by the basal ganglia, influenced by altered inputs from the cerebellum adapting to the FF. This interaction suggests potential interference originating from the interconnection between the basal ganglia and cerebellum, particularly if controller selection and motor adaptation are mediated by these two structures, respectively.

Another candidate locus of interference, suggested by the latency of ∼100ms associated with the exploitation of target redundancy, is the pathway linking the visuomotor system to the sensorimotor network through the parietal cortex. Omrani et al. (2016) showed that parietal area A5 modulated control responses during task engagement. This region is also involved in visuomotor processing for limb motor actions (Kalaska, 1996). It could thus signal the change in target shape and mediate the subsequent change in control policy. Additionally, its role in proprioceptive limb position estimate (Rushworth et al., 1997) makes it a likely candidate for adaptation to a new environment in which the forces applied to the arm vary. However, further investigation is needed to understand the role of each of the structures involved in these motor control mechanisms and confirm the origin of the observed interference. Importantly, the timing of control updates, as well as the indication that interference occurred when these changes happened during movement, provide temporal constraints for future work on the computational and neural bases of these processes.

In all, this study introduced a novel paradigm for examining the interplay between task-dependent control and sensorimotor adaptation. This new approach allowed us to better understand how these two mechanisms interact and might help to develop more refined methods for studying their relationship in healthy and clinical populations.

## Author Contributions

AD, PL and FC designed research; AD performed research; AD, PL an FC analyzed data; AD, PL and FC wrote the paper.

## Acknowledgements

A. D. was supported by Fonds de la Recherche Scientifique (F.R.S.-FNRS P.D.R. – Projet de Recherche). F. C. was supported by the F.R.S.-FNRS Grant 1.C.033.18. This work was additionally supported by a Concerted Research Action of Université catholique de Louvain (ARC; “coAction”) and from the WEL Research Institute (WELBIO Advanced Grant).

## Bibliography

1. Ahmadi-Pajouh MA., Towhidkhah F, Shadmehr R (2012) Preparing to Reach: Selecting an Adaptive Long-Latency Feedback Controller. J Neurosci 32(28):9537–9545

2. Bostan AC, Strick PL (2018) The basal ganglia and the cerebellum: nodes in an integrated network. Nat Rev Neurosci 19:338–350

3. Burdet E, Osu R, Franklin DW, Milner TE, Kawato M (2001) The central nervous system stabilizes unstable dynamics by learning optimal impedance. Nature 414(6862):446–9

4. Carey MR (2024) The cerebellum. Curr Biol 34(1), R7–R11.

5. Česonis J, Franklin DW (2022) Contextual cues are not unique for motor learning: Task-dependent switching of feedback controllers. PLoS Comput Biol 18(6):e1010192

6. Cluff T, Scott SH (2013) Rapid Feedback Responses Correlate with Reach Adaptation and Properties of Novel Upper Limb Loads. J Neurosci 33(40):15903–15914

7. Coltman S, Gribble P (2020) Time course of changes in the long-latency feedback response parallels the fast process of short-term motor adaptation. J. Neurophysiol 124(2):388–399

8. Crevecoeur F, Kurtzer I, Bourke T, Scott SH (2013) Feedback responses rapidly scale with the urgency to correct for external perturbations. J Neurophysiol 110(6):1323–1332.

9. Crevecoeur F, Scott SH, Cluff T (2019) Robust Control in Human Reaching Movements: A Model-Free Strategy to Compensate for Unpredictable Disturbances. J Neurosci 39(41):8135–8148

10. Crevecoeur F, Mathew J, Bastin M, Lefèvre P (2020a) Feedback Adaptation to Unpredictable Force Fields in 250 ms. ENeuro, 7(2)

11. Crevecoeur F, Thonnard J-L, Lefèvre P (2020b) A Very Fast Time Scale of Human Motor Adaptation: Within Movement Adjustments of Internal Representations during Reaching. Eneuro, 7(1)

12. Cross KP, Cluff T, Takei T, Scott SH (2019) Visual Feedback Processing of the Limb Involves Two Distinct Phases. J Neurosci 39(34):6751–6765

13. De Comite A, Crevecoeur F, Lefèvre P (2021) Online modification of goal-directed control in human reaching movements. J Neurophysiol 125(5):1883–1898.

14. Dizio P, Lackner JR (1995) Motor Adaptation to Coriolis Force Perturbations of Reaching Movements: Endpoint but not Trajectory Adaptation Transfers to the Nonexposed Arm. J Neurophysiol 74(4):1787–1792

15. Franklin DW, Wolpert DM (2008) Specificity of Reflex Adaptation for Task-Relevant Variability. J Neurosci 28(52):14165–14175

16. Franklin S, Wolpert DM, David X, Franklin W (2017) Rapid visuomotor feedback gains are tuned to the task dynamics. J Neurophysiol 118:2711–2726

17. Heald JB, Lengyel M, Wolpert DM (2021) Contextual inference underlies the learning of sensorimotor repertoires Nature 600(7889):489–493

18. Izawa J, Shadmehr R (2008) On-Line Processing of Uncertain Information in Visuomotor Control. J Neurosci 28(44):11360–11368

19. Joiner WM, Sing GC, Smith, MA (2017) Temporal specificity of the initial adaptive response in motor adaptation. PLoS Comput Biol 13(7): e1005438

20. Kalaska JF (1996) Parietal cortex area 5 and visuomotor behavior. Can J Physiol Pharmacol 74(4):483–498

21. Kalidindi HT, Crevecoeur F (2023) Human reaching control in dynamic environments. Curr Opin Neurobiol 83:102810

22. Kording KP, Tenenbaum JB, Shadmehr R (2007) The dynamics of memory as a consequence of optimal adaptation to a changing body. Nat Neurosci 10(6):779–786

23. Kurtzer IL, Pruszynski JA, Scott SH (2008) Long-latency reflexes of the human arm reflect an internal model of limb dynamics. Curr Biol 18(6):449–453

24. Kurtzer I, Trautman P, Rasquinha RJ, Bhanpuri NH, Scott SH, Bastian AJ (2013) Cerebellar damage diminishes long-latency responses to multijoint perturbations. J Neurophysiol 109:2228–2241

25. Lackner JR, DiZio P (2005) Motor control and learning in altered dynamic environments. Curr Opin Neurobiol 15(6):653–659

26. Liu D, Todorov E (2007) Evidence for the flexible sensorimotor strategies predicted by optimal feedback control. J Neurosci 27(35):9354–9368

27. Maeda RS, Gribble PL, Pruszynski JA (2020) Learning New Feedforward Motor Commands Based on Feedback Responses. Curr Biol 30(10):1941–1948

28. Murphy KP (2012) Machine learning: A probabilistic perspective. MIT Press.

29. Nashed JY, Crevecoeur F, Scott SH (2012) Influence of the behavioral goal and environmental obstacles on rapid feedback responses. J Neurophysiol 108(4):999–1009

30. Omrani M, Murnaghan CD, Pruszynski JA, Scott SH (2016) Distributed task-specific processing of somatosensory feedback for voluntary motor control. eLife 5:e13141

31. Orban de Xivry JJ, Lefèvre P (2016) A switching cost for motor planning. J Neurophysiol 116(6):2857–2868

32. Pruszynski JA, Kurtzer I, Lillicrap TP, Scott SH (2009) Temporal evolution of “automatic gain-scaling.” J Neurophysiol 102(2):992–1003

33. Rushworth MFS, Nixon PD, Passingham RE (1997) Parietal cortex and movement. I. Movement selection and reaching. Exp Brain Res 117(2):292–310

34. Scott SH (2004) Optimal feedback control and the neural basis of volitional motor control. Nat Rev Neurosci 5(7):532–544

35. Shadmehr R, Krakauer JW (2008) A computational neuroanatomy for motor control. Exp Brain Res 185(3):359–381

36. Shadmehr R, Mussa-Ivaldi FA (1994) Adaptive representation of dynamics during learning of a motor task. J Neurosci 14(5):3208–3224

37. Shadmehr R, Smith MA, Krakauer JW (2010) Error correction, sensory prediction, and adaptation in motor control. Annu Rev Neurosci 33:89–108

38. Singh K, Scott SH (2003) A motor learning strategy reflects neural circuitry for limb control. Nat Neurosci, 6(4):399–403

39. Smith MA, Ghazizadeh A, Shadmehr R (2006) Interacting adaptive processes with different timescales underlie short-term motor learning. PLoS Biol 4(6):e179

40. Spampinato DA, Block HJ, Celnik PA (2017) Cerebellar–M1 connectivity changes associated with motor learning are somatotopic specific. J Neurosci 37(9):2377–2386

41. Strick PL (1983) The influence of motor preparation on the response of cerebellar neurons to limb displacements. J Neurosci 3(10):2007–2020

42. Todorov E, Jordan MI (2002a) A Minimal Intervention Principle for Coordinated Movement. Advances in Neural Information Processing Systems 15:27–34

43. Todorov E, Jordan MI (2002b) Optimal feedback control as a theory of motor coordination. Nat Neurosci 5(11):1226–1235.

44. Vaswani PA, Shmuelof L, Haith AM, Delnicki RJ, Huang VS, Mazzoni P, Shadmehr R, Krakauer JW (2015) Persistent Residual Errors in Motor Adaptation Tasks: Reversion to Baseline and Exploratory Escape. J Neurosci 35(17):6969–6977

45. Wei K, Körding K (2010) Uncertainty of feedback and state estimation determines the speed of motor adaptation. Front Comput Neurosci 4:11

